## Supplementary PDF for "Scrutinizing the protein hydration shell from molecular dynamics simulations against consensus small-angle scattering data"

### Supplementary Material Table of Contents


**Supplementary Discussion**

Choice of SAS data used for quantitative comparison with MD simulations 3

Validation with $R_g$ from $P(r)$ vs. Guinier analysis 4

**Supplementary Methods: additional control SAS calculations**

Counter ion cloud contributions outside the envelope are negligible 4

Effect of salt relative to using only counter ions 5

Effect of cutoff settings for Lennard-Jones interactions 6

Effect of refined Lennard-Jones parameters to avoid $Na^+$–carboxylate overbinding in AMBER simulations (CUFIX) 6

Density correction for accurate buffer matching is critical for $R_g$ predictions 7

**Supplementary Figures**

S1 Visual representations of five proteins 9

S2 Schematic representation of hydration shell contrast for SAXS, SANS/$H_2O$, SANS/$D_2O$ and effect on $\Delta R_g$ 10

S3 Effect of protein flexibility 11

S4 Absolute $R_g$ values of five proteins 12

S5 $\Delta R_g^{SAS}$ values from simulations with position restraints on backbone atoms 13

S6 RNaseA structures from Amber99SBws–OPC simulations 14

S7 $\Delta R_g^{SAS}$ for lysozyme and urate oxidase 15

S8 $\Delta R_g^{SAS}$ compared to experimental $R_g$ values from Guinier analysis 16

S9 Effects of salt vs. only counter ions for xylanase 17

S10 Effects of salt vs. only counter ions for RNaseA 18

S11 Effects of salt vs. only counter ions for glucose isomerase 19

S12 $\Delta R_g^{SAS}$ values for glucose isomerase from simulations using only counter ions for 18 force field combinations 20

S13 Modified cation–carboxylate Lennard-Jones parameters with Amber force fields (CUFIX) 21

S14 Force-switch vs. plain cutoff with CHARMM36m 22

**Supplementary Tables**

S1 List of force fields and water models considered in this study 23

S2 Experimental consensus $R_g$ values from Trewhella et al.[1] 24

S3 All computed $\Delta R_g$ values of RNaseA 25

S4 All computed $\Delta R_g$ values of xylanase 26

S5 All computed $\Delta R_g$ values of glucose isomerase 27

S6 All computed $\Delta R_g$ values of lysozyme 28

S7 All computed $\Delta R_g$ values of urate oxidase 29

#### Supplementary Discussion

##### Choice of SAS data used for quantitative comparison with MD simulations

Trewhella, Vachette, *et al.* recently reported a worldwide round-robin benchmark on the reproducibility of SAXS and SANS experiments, involving experiments at 12 SAXS and four SANS instruments and leading to a total of 171 SAXS and 76 SANS curves on the following five proteins (Fig. S1): ribonuclease A (RNaseA, 13.690 kDa), hen egg white lysozyme (14.313 kDa), xylanase (20.844 kDa), urate oxidase (136.303 kDa), and glucose isomerase (172.910 kDa, also referred to as xylose isomerase).[1] Both, SAXS and SANS data were collected in batch mode or coupled to size-exclusion chromatography (SEC-SAXS, SEC-SANS). The $R_\mathrm{g}$ values were evaluated using Guinier analysis and based on the $P(r)$ function. Generally, good agreement was found between the experimental SAS curves and explicit-solvent SAXS predictions in the round-robin study.[1]

However, the quality and reproducibility of the SAS data differed among the five proteins. Specifically, $R_\mathrm{g}$ values for lysozyme varied substantially between different data sets, even if experiments were carried out in SEC mode. These variations have been attributed to an increased sensitivity of lysozyme to radiation damage[1] and are in line with variations of the lysozyme $R_\mathrm{g}$ in earlier reports.[2,3]

Urate oxidase exhibits increased propensity for aggregation and, thus, had to be shipped on ice and with a reduced concentration of only $5\,\mathrm{mg\,ml^{-1}}$.[1] In consequence, SAS data of urate oxidate was subject to poorer statistics as compared to the data of the other four proteins, in particular at wider scattering angles. Because the SAS curve over the whole $q$-range is used when computing $R_\mathrm{g}$ via the $P(r)$ function, such problems may lead to increased uncertainties of the $R_\mathrm{g}$ estimate. Furthermore, SANS in $H_2O$ exhibited exceptionally high background, and yielded unrealistic $R_\mathrm{g}$ values from Guinier analysis. Owing to these problems with lysozyme and urate oxidase, we did not include these proteins in the quantitative comparison of $\Delta R_\mathrm{g}^\mathrm{SAS}$ values in Fig. ??. For the sake of completeness, the $R_\mathrm{g}^\mathrm{SAS}$ values for lysozyme and urate oxidase, together with the experimental $R_{\mathrm{g}}$ estimates from Guinier analysis and $P(r)$, are shown in Fig. S7.

##### Validation with $R_{\mathrm{g}}$ from $P(r)$ vs. Guinier analysis

In this study, we used the experimental $R_{\mathrm{g}}$ values obtained from the $P(r)$ function and not from the Guinier analysis for comparison with MD simulations. In contrast to the Guinier analysis that uses only the small-angle region of the SAS curve, the $P(r)$ function makes use of the entire SAS curve. Thus, $R_{\mathrm{g}}$ values from the $P(r)$ reported in the round-robin SAS benchmark study were less sensitive to remaining undetected protein–protein aggregation and subject to smaller statistical uncertainties as compared to $R_{\mathrm{g}}$ estimates from the Guinier analysis (Table S2).[1] Nevertheless, the $\Delta R_{\mathrm{g}}^{\mathrm{SAS}}$ values from simulations reveal reasonable agreement also with $R_{\mathrm{g}}$ from Guinier analysis (Fig. S8).

#### Supplementary Methods: additional control SAS calculations

##### Counter ion cloud contributions outside the envelope are negligible

For the highly anionic glucose isomerase (GI), the counter ion cloud adds to the contrast of the hydration layer. According to Debye-Hückel theory, the counter ion cloud decays with the Debye length $\lambda_D$ into the bulk solvent, which equals $\lambda_D = 9.7\,\text{Å}$ for a 100 mM NaCl solution. Hence, since only solvent atoms within the envelope at a distance of 9 Å from the protein (Fig. ??A, blue surface) contributed to our explicit-solvent SAS calculations, counter ion cloud effects were likewise only taken into account up to a distance of 9Å from the protein surface, whereas counter ion cloud contributions beyond 9Å were neglected.

To exclude that this approximation affects the computed SAS curves, we estimated the effect of the counter ion cloud at distances beyond 9 Å on $R_{\mathrm{g}}$. Following Ref. 4, we modeled GI as a sphere with contrast, volume, and $R_{\mathrm{g}}$ taken from GI, and we used linearized Poisson–Boltzmann calculations to obtain the number densities of sodium and chloride as function of distance $R$ from the sphere surface. By computing $R_{\mathrm{g}}$ including the counter ion cloud up to $R = 9\,\text{Å}$ or up to a large distance, we found that $R_\text{g}$ was modified by the cutoff at $R = 9\,\text{Å}$ by only $0.033\,\text{Å}$, $-0.062\,\text{Å}$ and $-0.03\,\text{Å}$ for SAXS, SANS in $H_2O$, and SANS in $D_2O$. Since these errors are within the statistical errors, these effects were neglected for further analysis.

##### Effect of salt relative to using only counter ions

Since GI carries a large negative charge ($-60\,e$), GI is surrounded by a counter ion cloud. The ion densities in the counter ion cloud decay into the bulk solvent with the Debye length $\lambda_D$, which is inversely proportional to the square root of the ion concentration in bulk solvent.[5] Thus, upon simulating with a smaller ion concentration relative to experimental conditions, the spatial extend of the counter ion cloud in simulation would exceed the experimental conditions; consequently, the counter ion cloud in simulations would, relative to the experiment, impose an electron density contrast at larger distances from the protein, which may lead to an overestimated $R_\text{g}$. We tested the effect of the ion concentration for GI by computing $R_\text{g}$, $\Delta R_\text{g}$, and $\Delta R_\text{g}^\text{SAS}$ values from simulations with only counter ions, 100 mM NaCl salt, or 150 mM NaCl salt (Fig. S11). We find that, with increasing salt concentration, $\Delta R_\text{g}^\text{SAS}$ values decrease, as expected from a deceasing Debye length and, thereby, from a spatially more compact counter ion cloud. For instance, in simulations with CHARMM36m–TIP3P, upon replacing counter ions with 150 mM NaCl salt, the $R_\text{g}$ from SAXS relative to SANS/$D_2O$ decreases by ∼0.2Å. This trend is confirmed by GI simulations with all 18 force field combinations using purely counter ions (compare Fig. S12 with Fig. **??**E/F). Thus, for quantitative comparison of $\Delta R_\text{g}^\text{SAS}$ values between simulation and experiment, an at least approximate match of the buffer conditions is mandatory, as used for our study.

In addition, we tested the effect of adding additional 150 mM NaCl to simulation of xylanase and RNaseA instead of using only counter ions (Figs. S9 and S10). For these near-neutral proteins, $\Delta R_\text{g}^\text{SAS}$ values agreed within error bars between simulations with salt and simulations with counter ions, as expected form the absence of a pronounced counter ion cloud.

##### Effect of cutoff settings for Lennard-Jones interactions

Neglect of Lennard-Jones (LJ) interactions beyond the cutoff could in principle modify the density of the hydration shell. Thus, we tested the effects of LJ cutoffs for xylanase simulated with the CHARMM36m–TIP3P force field combination. The CHARMM36m force field has been parameterized with a gradual switch of LJ forces between 1 and 1.2 nm. We carried out xylanase simulations either (i) following the CHARMM recommendations ("force-switch") or (ii) using a plain cutoff at 1 nm. For each cutoff setting, simulations were carried out with restraints on heavy atoms, restraints on the backbone, or using a free MD simulation (Fig. S14). For simulations with restraints on the backbone or for free MD simulations, we obtained larger $\Delta R_{\mathrm{g}}^{\mathrm{SAS}}$ values with the force-switch settings as compared to using a plain cutoff (Fig. S14, orange vs. yellow and lightblue vs. darkblue bars). Hence, upon validating force fields against $\Delta R_{\mathrm{g}}^{\mathrm{SAS}}$ values from experiments, it is critical to follow the cutoff settings that have been used during force field parametrization.

Notably, for simulations with restraints on heavy atoms, no such effect was visible (Fig. S14, pink vs. red bars). Thus, additional LJ interactions between 1 nm and 1.2 nm lead to larger $\Delta R_{\mathrm{g}}^{\mathrm{SAS}}$ only in the presence of amino acid side chain fluctuations. These findings demonstrate that the additional LJ interactions between 1 nm and 1.2 nm with force-switch settings lead to increased $\Delta R_{\mathrm{g}}^{\mathrm{SAS}}$ values not because of tighter packing of water on a given protein surface; instead, the additional interactions lead to more favorable protein–water conformations, which manifest in a larger density contrast of the hydration shell.

##### Effect of refined Lennard-Jones parameters to avoid $Na^+$–carboxylate overbinding in AMBER simulations (CUFIX)

Certain older models of monovalent cations such as $Na^+$ tend to overbind to carboxylate moieties, as present in aspartate and glutamate residues.[6,7] We found previously that such overbinding of monovalent ions to anionic proteins may influence the $R_g$ values obtained from explicit-solvent SAS calculations.[8] Whereas refined LJ parameters have been incorporated into the default CHARMM36m force field to avoid overbinding (denoted "NBFIX" or "CUFIX"), refined LJ parameters for AMBER force fields have become available only recently.[7]

As discussed in the Results, $\Delta R_g^{SAS}$ values for GI obtained with CHARMM36m were systematically larger as compared to values obtained with AMBER force fields, irrespective of the applied water model (Fig. **??**E/F). We hypothesized that different binding properties of $Na^+$ ions in CHARMM36m as compared to AMBER simulations may partly explain such effect. To test this hypothesis, we carried out additional SAS calculations for GI with the ff14SB force field with CUFIX and compared the $\Delta R_g^{SAS}$ values with results from standard ff14SB or CHARMM36m (Fig. S13).[7] Upon modifying the LJ interaction with CUFIX in ff14SB simulations, (i) the density of $Na^+$ at the protein surface decreased relative to ff14SB and took lower densities as compared to ff99SB-ildn (Fig. S13G), (ii) $\Delta R_g^{SAS}$ values increased by ~0.2Å or ~0.1Å for SANS/$D_2O$ or SANS/$H_2O$ (Fig. S13B/C, pink vs. orange bars). However, the $\Delta R_g^{SAS}$ values with ff14SB/CUFIX remained below the results from CHARMM36m. Thus, refined $Na^+$–carboxylate Lennard-Jones interactions have only a small effect on $\Delta R_g^{SAS}$ values, and the increased $\Delta R_g^{SAS}$ value with CHARMM36m relative to several AMBER force fields is only partly explained by reduced binding of $Na^+$ ions to the protein surface.

##### Density correction for accurate buffer matching is critical for $R_g$ predictions

Explicit-solvent SAS predictions implemented by GROMACS-SWAXS use a density correction for two reasons:[9,10] (i) Certain water models such as TIP3P underestimate the water density, which would lead to an overestimation of the contrast of the protein relative to bulk solvent and, thereby, to an overestimation of the forward scattering $I_0$. (ii) Since the bulk solvent density in the protein simulation may marginally differ from the density in the pure-solvent simulations, possibly owing to finite-size effects, spurious contrasts may emerge at larger distances from the protein, which would lead to artifacts in the $R_\mathrm{g}$ estimate.

To avoid such artifacts, GROMACS-SWAXS corrects the solvent density both in the pure-solvent and in the protein simulation system. In the pure-solvent system, a uniform density is added to match a preselected bulk density $\rho^0_\mathrm{bulk}$, which may be taken as the experimental density of pure water of $334\,\mathrm{e\,nm^{-3}}$. In the protein simulation, the bulk density $\rho^\mathrm{MD}_\mathrm{bulk}$ is obtained by averaging over simulation frames within the volume *outside* of the envelope. A correction factor is obtained as the difference relative to $\rho^0_\mathrm{bulk}$ as $f = \rho^0_\mathrm{bulk}/\rho^\mathrm{MD}_\mathrm{bulk}$. Then the solvent density *inside* of the envelope (which contributes to the calculated SAS curve) is corrected by the factor $f$.

Here, we tested the effect of the density correction on $\Delta R_\mathrm{g}^\mathrm{SAS}$ values. To this end, we first replaced the preselected bulk density of $334\,\mathrm{e\,nm^{-3}}$ with the bulk density of the respective water model. Thereby, the protein–bulk contrast may not match the experimental value, however, accurate buffer matching between the protein and the pure-solvent simulations was still active. As expected, this led to a considerable change of the forward scattering $I_0$ for water models with inaccurate bulk densities such as TIP3P or cTIP3P; however, the $\Delta R_\mathrm{g}^\mathrm{SAS}$ revealed only a marginal change that was not visible in the bar plots.

Second, we fully disabled the density correction, thereby not ensuring accurate buffer matching any more. In these calculations, we observed changes of $\Delta R_\mathrm{g}^\mathrm{SAS}$ of SAXS relative to SANS/$D_2O$ by up to 0.7Å. In addition, the results revealed poor agreement with experiments. Thus, for accurate $\Delta R_\mathrm{g}^\mathrm{SAS}$ and for comparison with experiments, corrections for accurate buffer matching are mandatory, as used in the present study.

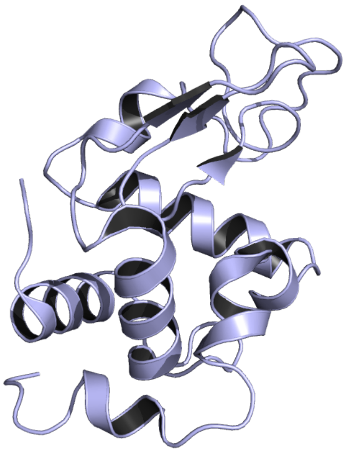

(A) Hen egg white lysozyme (PDB code 3L8W[11])

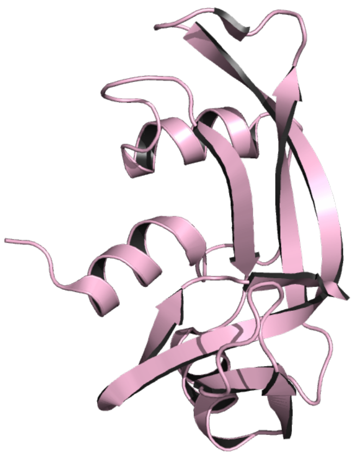

(B) Ribonuclease A (RNaseA; 7RSA[12])

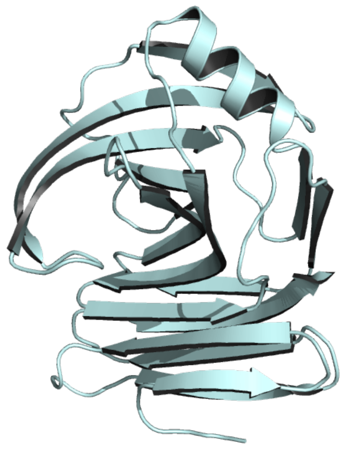

(C) Xylanase (2DFC[13])

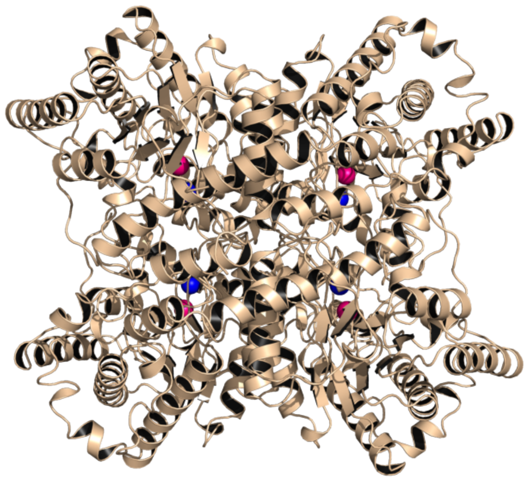

(D) Glucose isomerase. 1MNZ[14] after adding one methionine residue at the N- terminus. $Ca^{2+}$ and $Mg^{2+}$ ions inside the protein are shown as blue and pink spheres, respectively.

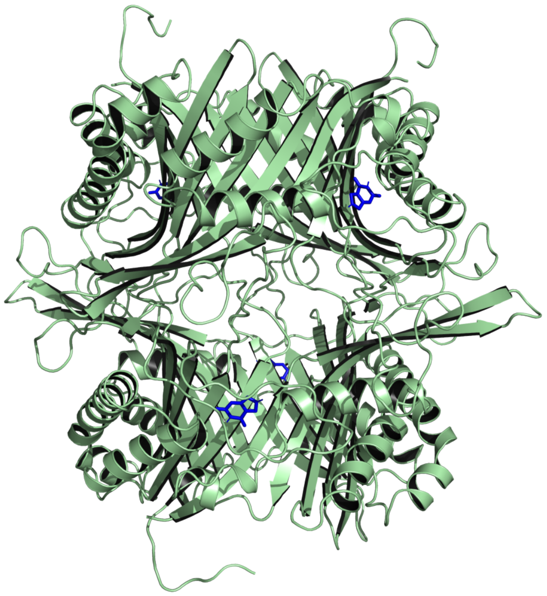

(E) Urate oxidase. 3L8W,[11] after adding six missing residues (sequence SLKSKL) at the C-terminus). Xanthine ligands are shown with blue sticks.

Figure S1: Cartoon representation of five proteins considered in this study (A) hen egg white lysozyme, (B) ribonuclease A (RNaseA), (C) xylanase , (D) glucose isomerase, and (E) urate oxidase.

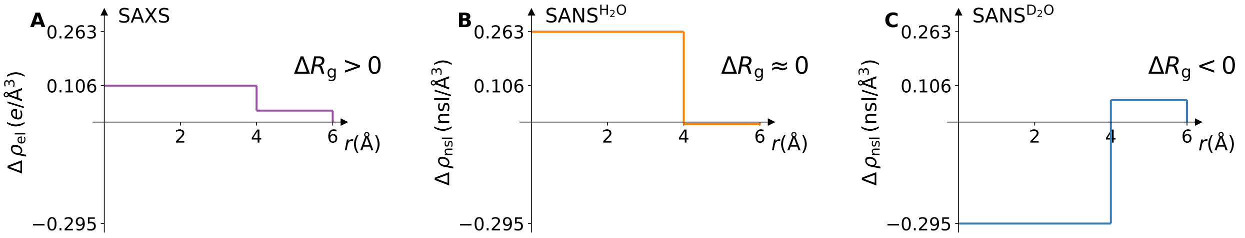


Figure S2: On the effect of the protein hydration shell on $R_g$ as detected by SAXS, SANS in $H_2O$, or SANS in $D_2O$, illustrated with a hypothetical spherical protein with a (unitless) radius of 4 and a thickness of the hydration shell of 2. (A–C) Scattering contrast relative to bulk as function of distance $r$ from the sphere center detected by (A) SAXS, (B) SANS/$H_2O$, and (C) SANS$D_2O$ experiments. (A) As detected by SAXS, both the protein and the hydration shell exhibit a positive electron density contrast relative to the bulk, resulting in an increased $R_g$ owing to the hydration shell ($\Delta R_g > 0$). (B) During SANS/$H_2O$, the protein exhibits a positive contrast of the neutron scattering length density, while the contrast of the hydration shell is close to zero, leading to a small influence by the hydration shell on $R_g$ ($\Delta R_g \approx 0$). (C) During SANS/$D_2O$, the protein exhibits a negative contrast whereas the hydration shell exhibits a positive contrast relative to bulk, resulting in a decreased $R_g$ ($\Delta R_g < 0$).

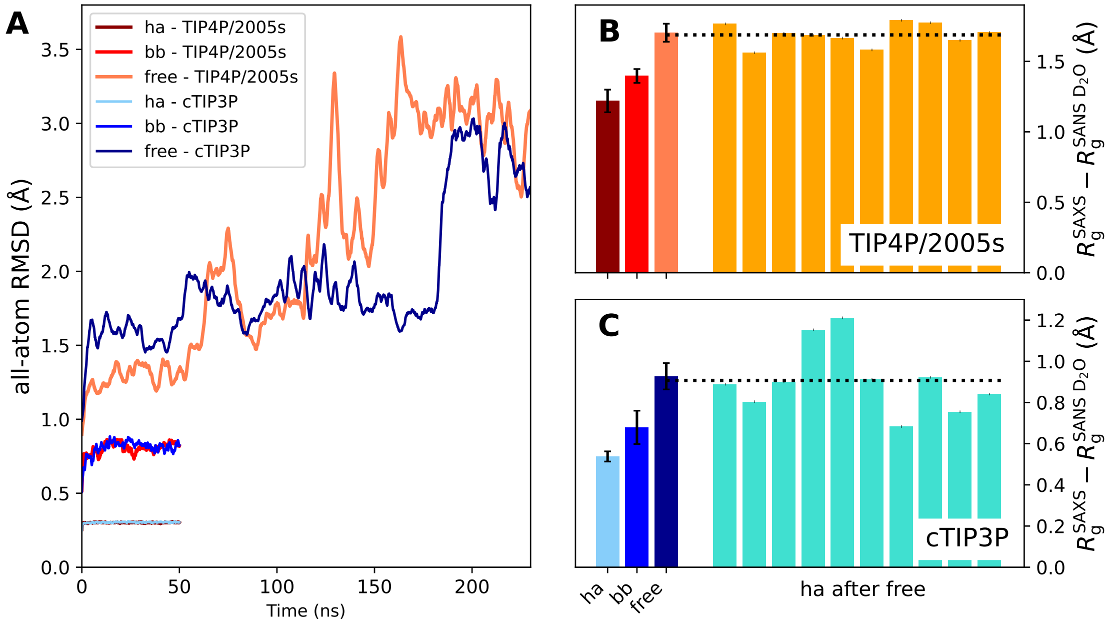


Figure S3: Effects of protein flexibility and hydration shell relaxation on $R_g$ in simulations of xylanase. (A) All-atom RMSD with ff99SBws–TIP4P/2005s (brown shades) or CHARMM36m–cTIP3P (blue shades) with increasing flexibility. Simulations were carried out with restraints on heavy atoms (ha, see legend), with restraints on the backbone (bb), or without restraints (free). (B) Difference $\Delta R_g^{SAS}$ between the $R_g$ from SAXS relative to SANS/$D_2O$ for amber99SBws–TIP4P/2005s and (C) CHARMM36m–cTIP3P. Left three bars: $\Delta R_g^{SAS}$ from simulations with restraints on heavy atoms, restraints on the backbone, or from free simulations starting from the crystal structure (color code according to panel A), revealing an increased $\Delta R_g^{SAS}$ with increasing protein flexibility. Right ten bars: $\Delta R_g^{SAS}$ values from MD simulations with restraints on all heavy atoms starting from different times of the free simulation. Critically, the average $\Delta R_g^{SAS}$ values from these ten restrained simulations (dotted lines) agree with $\Delta R_g^{SAS}$ from the initial free simulation. Thus, not the protein flexibility *per se*, but instead the relaxation of protein–water packing during the initial free MD leads on average to increased $\Delta R_g^{SAS}$ values.

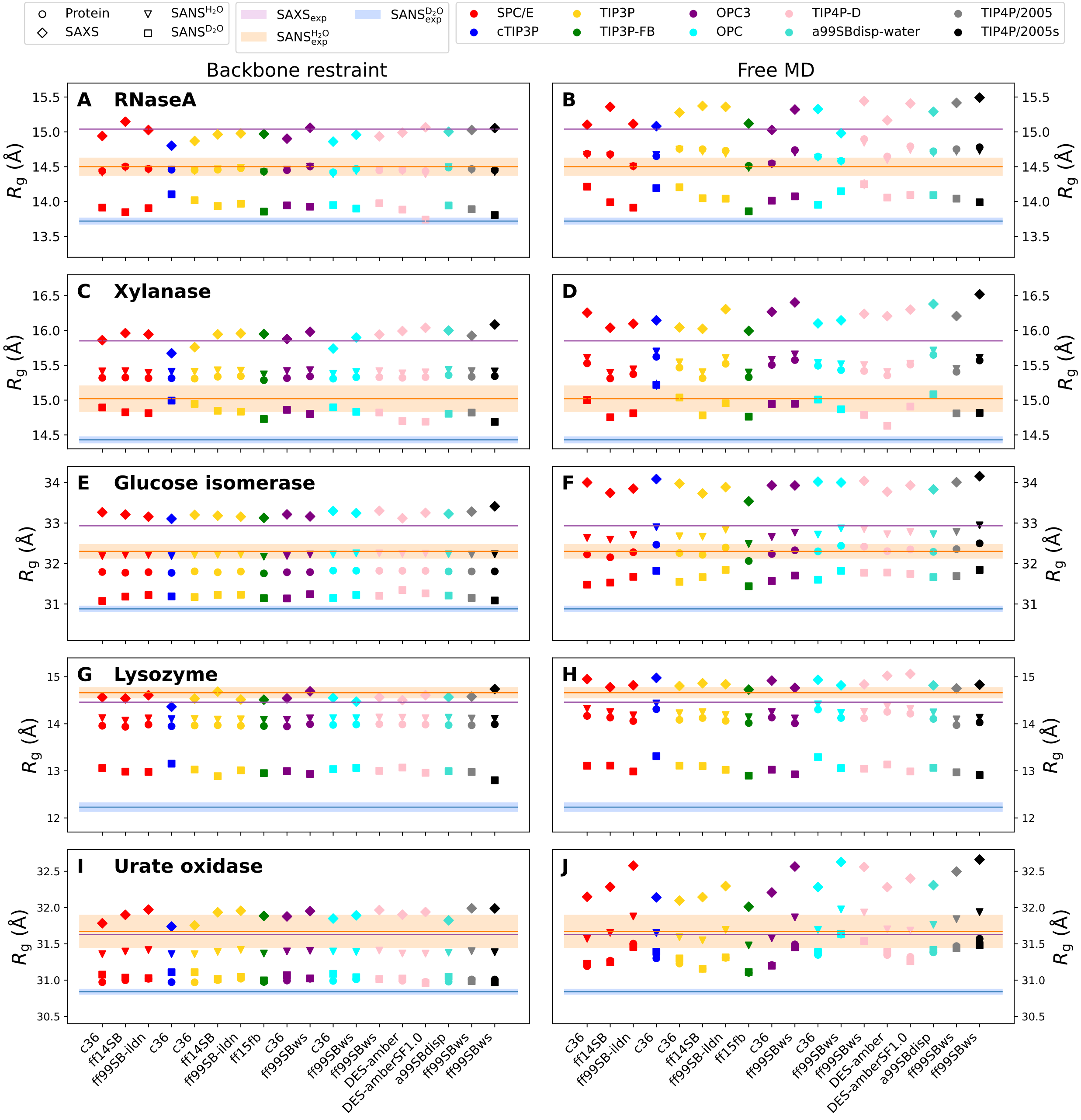


Figure S4: Absolute $R_\mathrm{g}$ values of (A/B) RNaseA, (C/D) xylanase, (E/F) glucose isomerase, (G/H) lysozyme, and (I/J) urate oxidase obtained from MD simulations with backbone restraints (left column) and without restraints (right column) with 18 different combinations of protein force fields (labels along the abscissa) and water models (color code, see legend). $R_g^{\mathrm{SAS}}$ values from free MD simulations for SAXS (diamonds), SANS/$H_2O$ (triangles), and SANS/$D_2O$ (squares). $R_\mathrm{g}$ of the pure protein ($R_g^{\mathrm{Prot}}$) shown as circles. Experimental $R_g^{\mathrm{SAS}}$ values from $P(r)$ analysis are shown has horizontal lines for SAXS (purple), SANS/$H_2O$ (orange), and SANS/$D_2O$ blue. Shaded areas indicated experimental uncertainties of consensus SAS data.[1] Absolute $R_\mathrm{g}$ values shown here are not suitable for validating the hydration shell against experimental data because $R_\mathrm{g}$ of the bare protein ($R_\mathrm{g}^{\mathrm{Prot}}$) is not accurately known in the experiment.

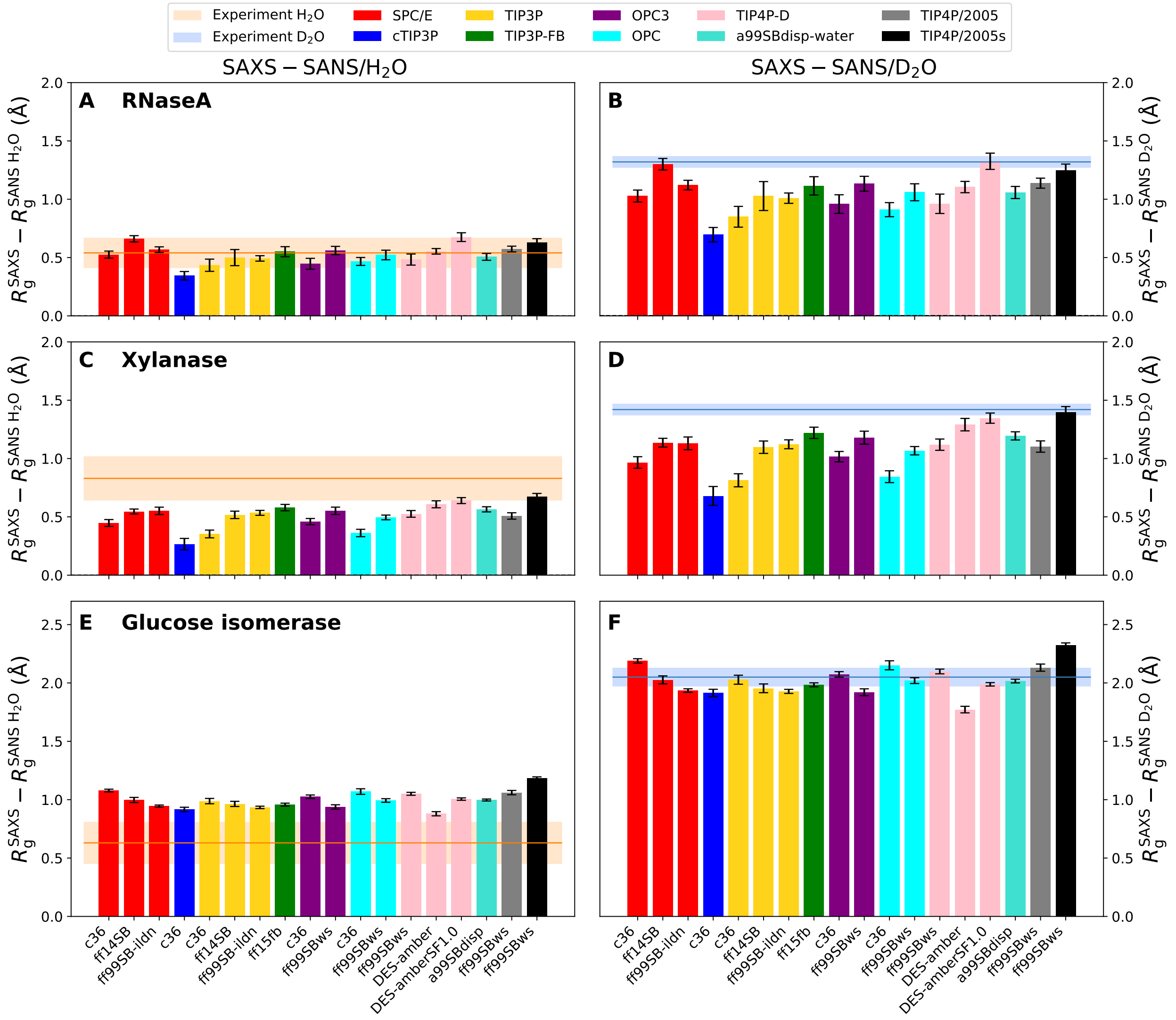


Figure S5: Same analysis as shown in Fig. ??, however based on simulations with backbone restraints instead of based on free simulations. Large variations of $\Delta R_g^{SAS}$ among different force field combinations are evident, even with identical backbone conformations. Thus, differences in protein–water interactions among different force fields and not different protein conformations in free simulations dominate the variations of $\Delta R_g^{SAS}$. As a second finding, $\Delta R_g^{SAS}$ values from backbone-restrained simulations (this figure) are mostly smaller as compared to values from free simulations (Fig. ??), demonstrating that the equilibration of protein–water interactions during free simulations leads to a (slightly) increased packing of the hydration shell (see also Fig. S3).

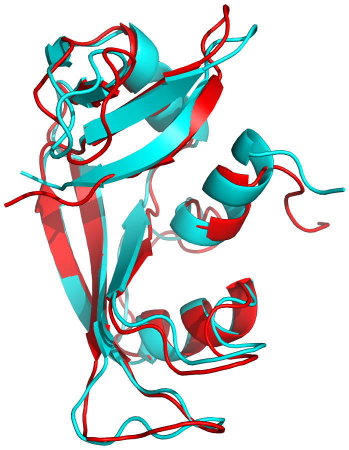

Figure S6: Structures of RNaseA in simulations with CHARMM36m–OPC after equilibration (turquoise) and at ∼100 ns of free MD simulation (red). The N-terminal helix partly unfolded in the free simulation. In simulation with other force field combinations, the N-terminal helix exhibited some flexibility as well, however, the helix re-folded during the simulations. The conformational instability of RNaseA in CHARMM36m–OPC simulations may explain the the large $\Delta R_{\mathrm{g}}^{\mathrm{SAS}}$ value in free simulations of RNAseA (Fig. **??**B, light turquoise bar labeled with "c36"), in contrast to all other free simulations with CHARMM36m (see Figs. **??**) or all restrained simulations CHARMM36m (including CHARMM36m–OPC, see Fig. S5).

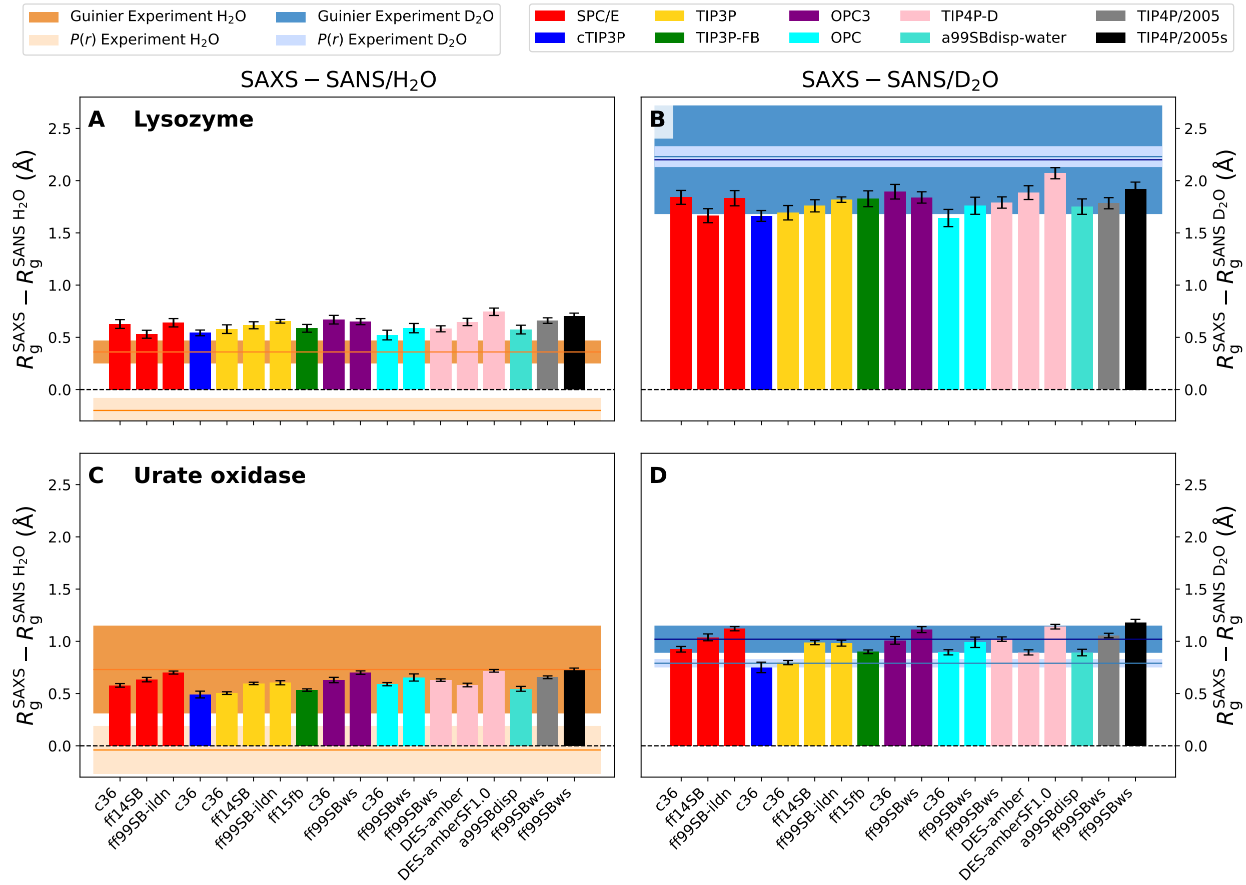


Figure S7: $\Delta R_{\mathrm{g}}^{\mathrm{SAS}}$ values of (A/B) lysozyme and (C/D) urate oxidase for SAXS relative to SANS/$H_2O$ (left column) and SAXS relative to SANS/$D_2O$ (right column). Experimental consensus data (horizontal lines) and reported uncertainties (shaded areas) from Guinier analysis (dark brown, dark blue) and from $P(r)$ analysis (light brown, light blue) are shown for reference. Experimental values for lysozyme and urate oxidase are subject to increased uncertainty, owing to problems with radiation damage and aggregation (see Supplementary Discussion).[1]

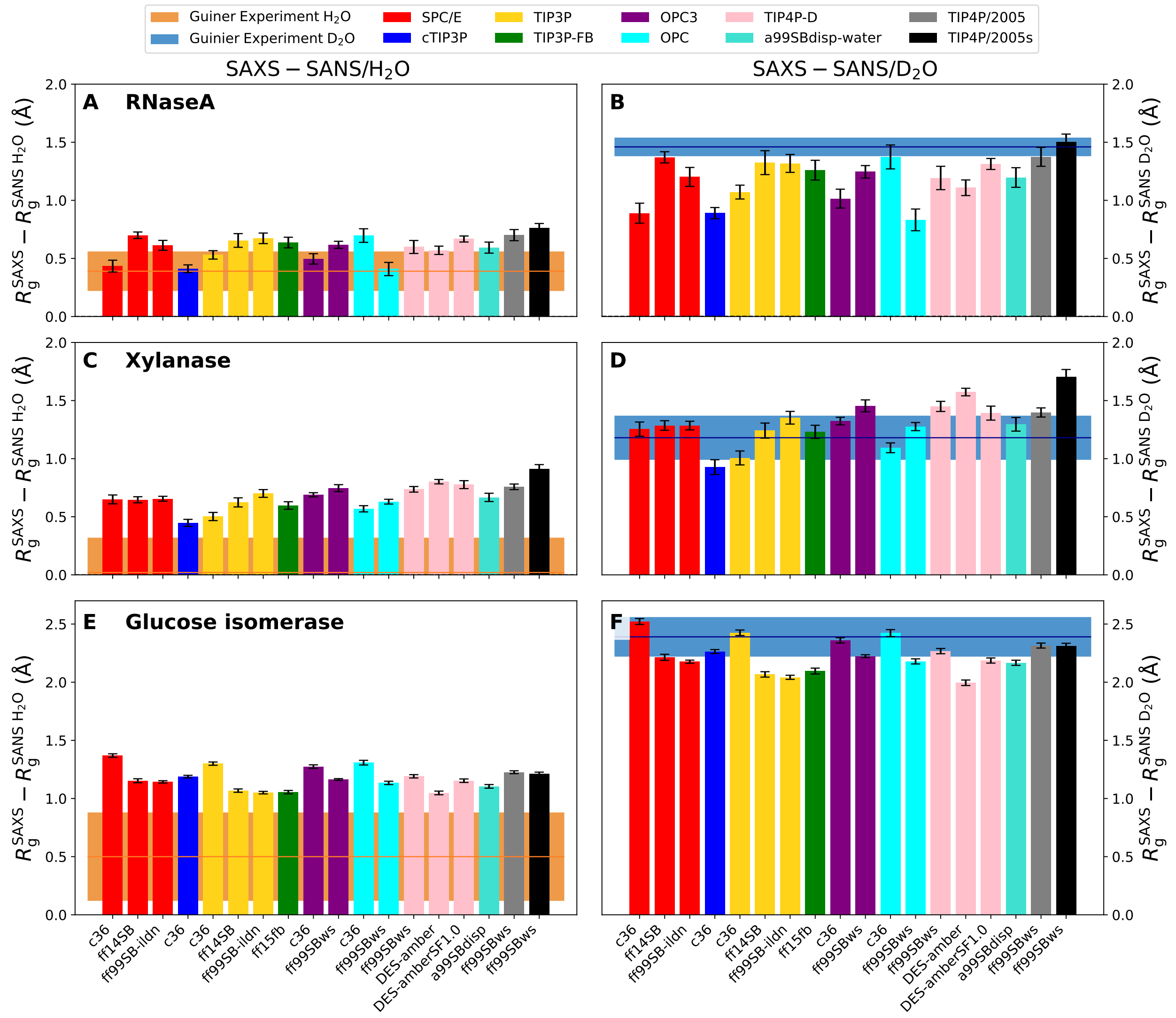


Figure S8: $\Delta R_g^{SAS}$ for (A/B) RNaseA, (C/D) xylanase, and (E/F) glucose isomerase, following the same labeling and color code as in Fig. ??. Experimental values obtained form Guinier analysis are shown as horizontal lines with uncertainties as shaded areas,[1] in contrast to experimental values from the $P(r)$ function in Fig. ??. The larger experimental errors as compared to the results from $P(r)$ analysis and the slightly poorer agreement between simulations and experiment (compared to Fig. ??) are rationalized by the fact that Guinier analysis is more sensitive to small amounts of undetected protein–protein aggregation.[1] Left column: $\Delta R_g^{SAS}$ from SAXS relative to SANS/$H_2O$ revealing poor agreement between simulation and experiment, likely caused by poorer signal-to-noise ratio during SANS/$H_2O$ experiments owing to increased incoherent scattering in $H_2O$ as compared to $D_2O$ and lower contrast.

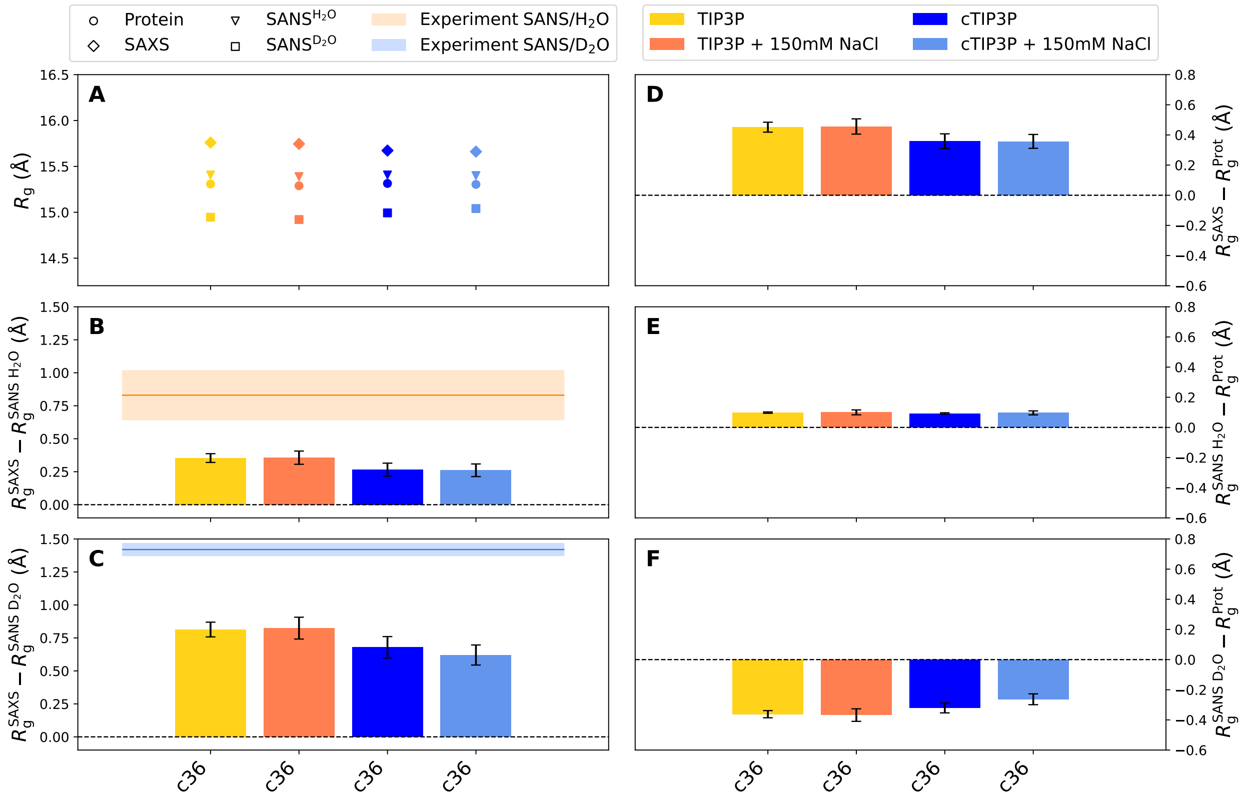


Figure S9: On the effect of 150 mM NaCl on SAS-derived $R_g$ values of xylanase as compared to simulations with purely counter ions. Simulations were carried out with Charmm36m combined with three different water models using purely counter ions (SPC/E, TIP3P, or cTIP3P, see legend for color code) or with 150 mM NaCl (denoted SPC/E-ions, TIP3P-ions, cTIP3P-ions, see legend). (A) SAS-derived $R_g^{SAS}$ computed from free MD simulations for SAXS (diamonds), SANS/$H_2O$ (triangles), and SANS/$D_2O$ (squares). $R_g$ of the pure protein ($R_g^{Prot}$) shown as circles. Experimental $R_g^{SAS}$ values from $P(r)$ analysis are shown has horizontal lines for SAXS (purple), SANS/$H_2O$ (orange), and SANS/$D_2O$ (blue). (B) Difference between the $R_g$ from SAXS relative to SANS/$H_2O$ and (C) between SAXS relative to SANS/$D_2O$. (D) Difference of SAS-derived $R_g$ from explicit-solvent MD relative to $R_g^{Prot}$ for SAXS,(E) for SANS/$H_2O$, and (F) for SANS/$D_2O$. Statistical errors (1 SE) were obtained from block averaging. Horizontal lines and shaded areas indicated experimental consensus values and uncertainties.[1] Adding 150 mM NaCl to the buffer has only a small effect on the $\Delta R_g$ values of the only slightly charged xylanase.

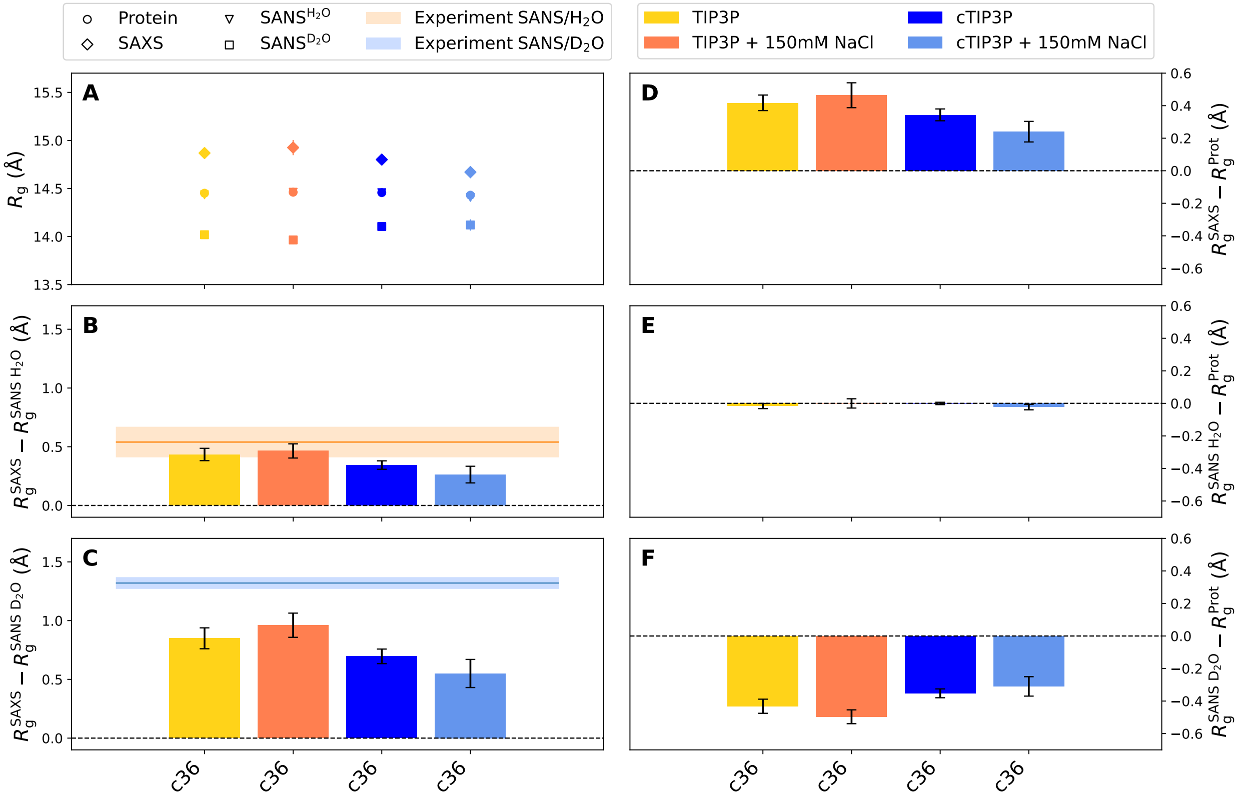


Figure S10: On the effect of 150 mM NaCl on SAS-derived $R_g$ values of RNaseA. Same analysis and as shown for xylanase in Fig. S9.

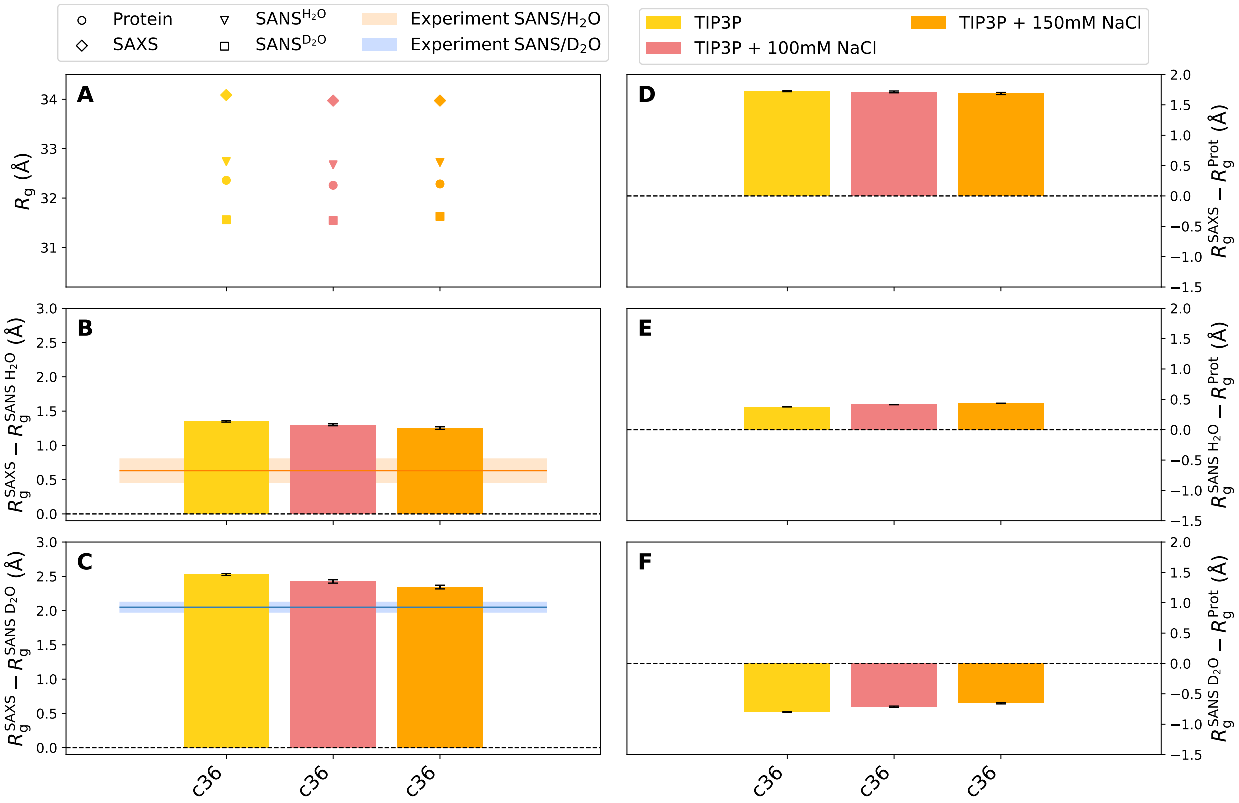


Figure S11: On the effect of 100 mM NaCl (pink) or 150 mM NaCl (orange) on SAS-derived $R_\mathrm{g}$ values of the highly anionic glucose isomerase ($-60\,e$), as compared to using only counter ions (yellow). Same analysis as shown for xylanase and RNaseA in Figs. S9 and S10, however restricted to TIP3P water.The lower the ion concentration in the buffer, the larger the Debye length and thus the effect of the hydration shell on the $R_\mathrm{g}$.

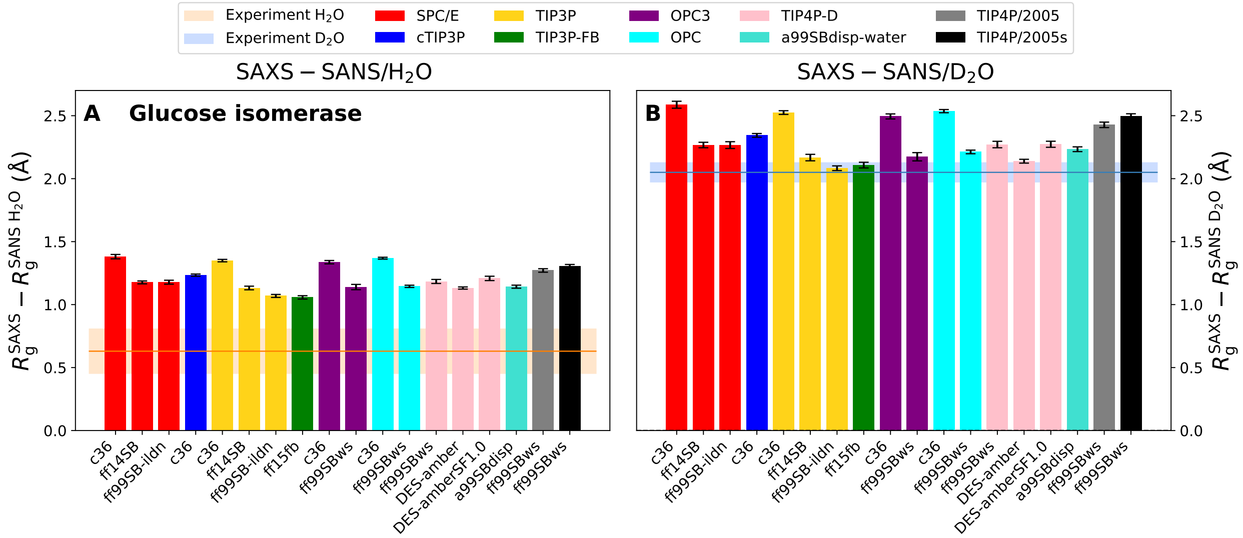


Figure S12: On the importance of salt for accurate $R_g$ predictions of highly anionic glucose isomerase. (A) $\Delta R_g^{SAS}$ values from SAXS relative to SANS/$H_2O$ and (B) of SAXS relative to SANS/$D_2O$ from free simulations of glucose isomerase with only counter ions (60 $Na^+$) but no additional salt. Experimental consensus values and uncertainties from $P(r)$ analysis are shown as horizontal lines and shaded areas, respectively. By comparison with Fig. ??E/F, the lack of salt increases $\Delta R_g^{SAS}$ systematically, caused by an overly extended counter ion cloud. In turn, in presence of salt, the Debye length decreases, thus leading to a spatially more compact counter ion cloud and slightly smaller $\Delta R_g^{SAS}$ values (compare with Fig. ??).

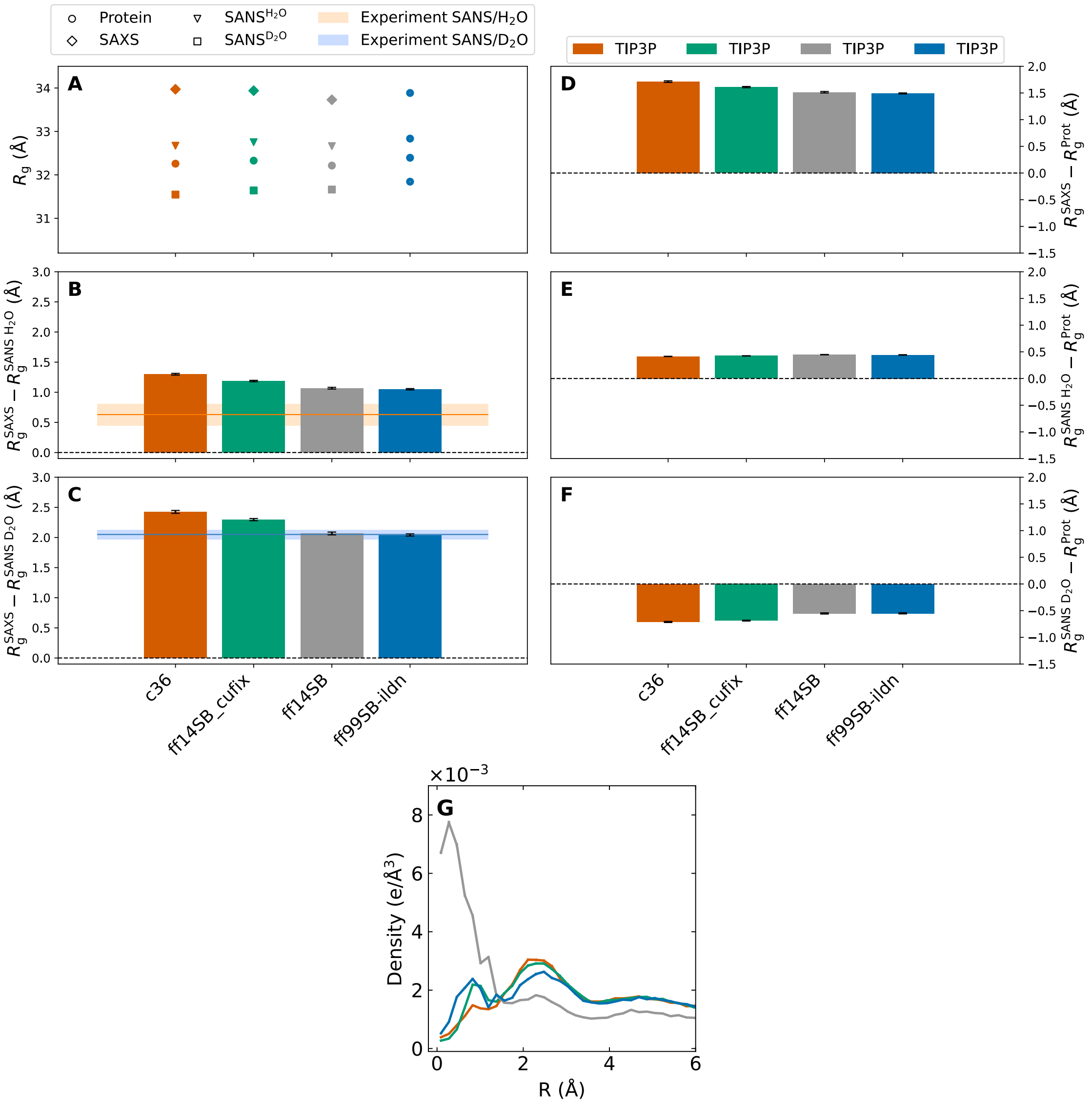


Figure S13: On the effect of non-bonded fix parameters (CUFIX, also called NBFIX) with the Amber14SB protein force field. CUFIX implements modified Lennard-Jones interactions between $Na^+$ ions and carboxylate oxygen atoms of aspartate and glutamate residues with the aim to avoid $Na^+$–$COO^-$ overbinding.[7] CHARMM36m implements CUFIX by default in the version of July 2020 as used here. (A) $R_g$, (B/C) $\Delta R_g^{SAS}$, and (D–F) $\Delta R_g$ values are reported for the highly anionic glucose isomerase (GI, $-60\,e$). Free simulations of GI were carried out with CHARMM36m (yellow bar), ff14SB with CUFIX (pink), as well as ff14SB or ff99SB-ildn without CUFIX (grey and blue, respectively). Experimental consensus values and uncertainties from $P(r)$ analysis are shown as horizontal lines and shaded areas, respectively. Statistical errors (1 SE) were obtained from block averaging. Upon refining $Na^+$–carboxylate interactions by ff14SB with CUFIX, $\Delta R_g^{SAS}$ increased by only ~0.2 Å and ~0.1Å for SANS in $D_2O$ and $H_2O$, respectively. (G) Density of $Na^+$ as function of distance from the protein surface for four different force fields (color code according to panels A–F). CUFIX greatly reduces $Na^+$ overbinding in ff14SB simulations.

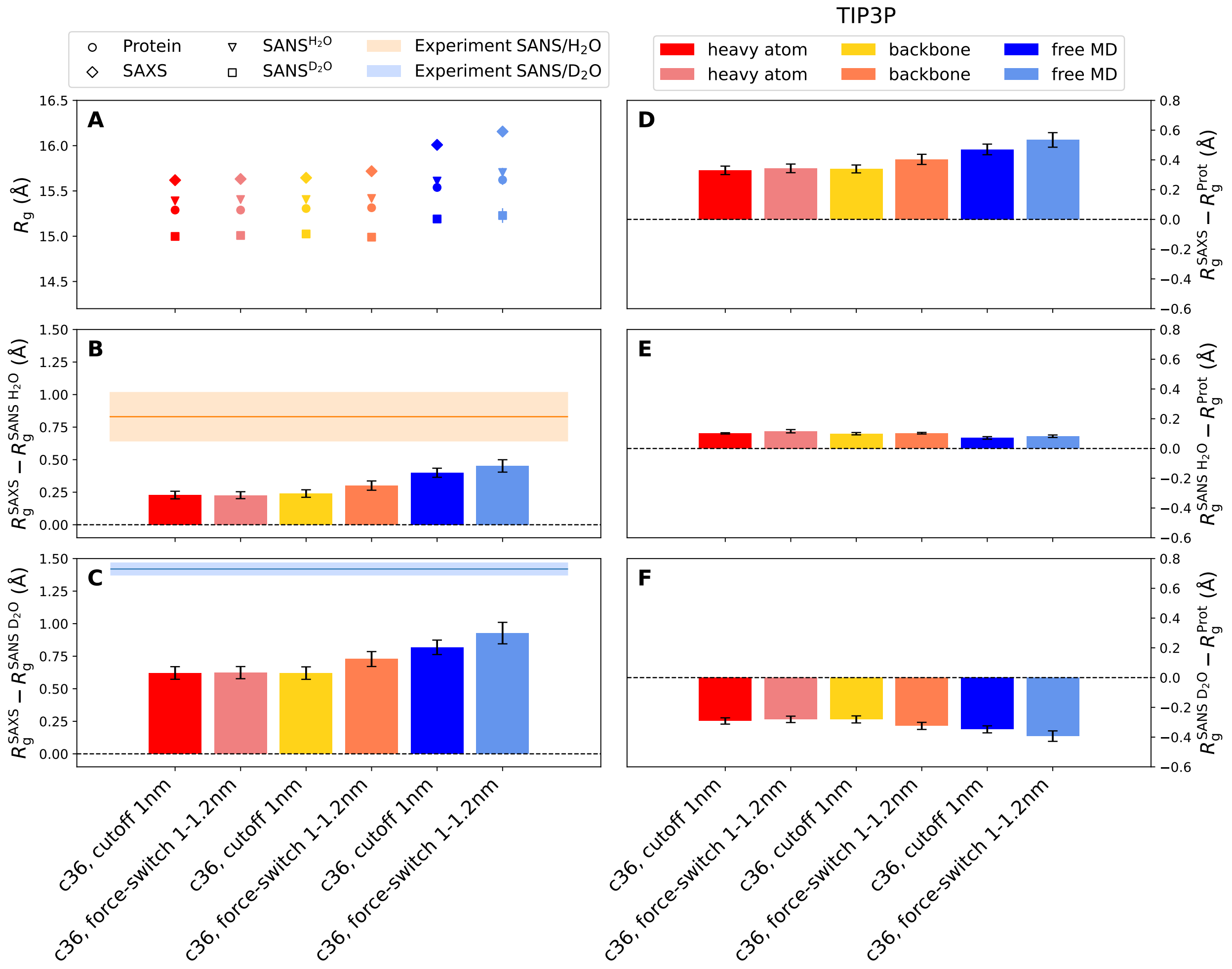


Figure S14: On the role of force field-specific Lennard-Jones (LJ) cutoff settings: (A) $R_g$, (B/C) $\Delta R_g^{SAS}$, and (D–F) $\Delta R_g$ values for xylanase using the CHARMM36m–TIP3P force field combination. Simulations were carried out either using a plain LJ cutoff at 1.0 nm or by switching off the LJ forces between 1 nm and 1.2 nm, as recommended for the CHARMM36m force field ("force-switch 1–1.2 nm"). Results are shown for simulations with restrained heavy atoms (red, pink), restrained backbone (yellow, orange) or for free MD (dark and light blue). In simulations with flexible side chains (free or backbone-restrained MD), cutoff settings following CHARMM standards leads to larger $\Delta R_g^{SAS}$ values as compared to using a plain 1 nm cutoff (panels B/C, orange vs. yellow, light blue vs. dark blue). In simulations with restrained heavy atoms, using a different cutoff stetting has only a marginal effect (pink vs. red). Thus, the additional LJ interactions between 1 nm and 1.2 nm lead to increased $\Delta R_g^{SAS}$ values owing to more densely packed water structures around flexible protein side chains, and not owing to stronger attraction of water onto a (fixed) protein surface.

Table S1: Eighteen combinations of protein force field and water model used in this study. CHARMM36m was taken from version of July 2020.

| Protein force field | Abbreviation | Water model | Refs. |
|---|---|---|---|
| CHARMM36m | c36 | cTIP3P | 15, 16 |
| | | TIP3P | 17 |
| | | SPC/E | 18 |
| | | OPC3 | 19 |
| | | OPC | 20 |
| Amber14SB | ff14SB | SPC/E | 21 , 18 |
| | | TIP3P | 17 |
| Amber99SB-ildn | ff99SB-ildn | SPC/E | 22, 18 |
| | | TIP3P | 17 |
| Amber15/force-balance | ff15fb | TIP3P-FB | 23 |
| Amber99SBws | ff99SBws | TIP4P/2005s | 24, 25 |
| | | TIP4P/2005 | 26 |
| | | TIP4P-D | 27 |
| | | OPC3 | 19 |
| | | OPC | 20 |
| DES-amber | DES-amber | TIP4P-D | 28, 27 |
| DES-amber without scaled charges | DES-amberSF1.0 | TIP4P-D | 28, 27 |
| Amberr99SBdisp | a99SBdisp | a99SBdisp-water | 29 |

Table S2: Experimental consensus $R_g$ values for SAXS, SANS/$H_2O$, and SANS/$D_2O$ from Trewhella *et al.*[1] $R_g$ are shown as obtained from Guinier or $P(r)$ analysis. Two right columns: $\Delta R_g^{SAS}$ values from SAXS relative to SANS/$H_2O$ and SAXS relative to SANS/$D_2O$, respectively.

| Protein | Type of $R_g$ | $R_g^{SAXS}$ (Å) | $R_g^{SANS\ H_2O}$ (Å) | $R_g^{SANS\ D_2O}$ (Å) | $R_g^{SAXS} - R_g^{SANS\ H_2O}$ (Å) | $R_g^{SAXS} - R_g^{SANS\ D_2O}$ (Å) |
|---|---|---|---|---|---|---|
| RNaseA | Guinier | 15.13 ± 0.02 | 14.74 ± 0.17 | 13.67 ± 0.08 | 0.39 ± 0.17 | 1.46 ± 0.08 |
| | $P(r)$ | 15.04 ± 0.01 | 14.50 ± 0.13 | 13.72 ± 0.05 | 0.54 ± 0.13 | 1.32 ± 0.05 |
| Xylanase | Guinier | 16.05 ± 0.01 | 16.03 ± 0.30 | 14.87 ± 0.19 | 0.02 ± 0.3 | 1.18 ± 0.19 |
| | $P(r)$ | 15.85 ± 0.01 | 15.02 ± 0.19 | 14.43 ± 0.05 | 0.83 ± 0.19 | 1.42 ± 0.05 |
| Glucose isomerase | Guinier | 33.11 ± 0.05 | 32.61 ± 0.38 | 30.72 ± 0.17 | 0.5 ± 0.38 | 2.39 ± 0.17 |
| | $P(r)$ | 32.93 ± 0.01 | 32.30 ± 0.18 | 30.88 ± 0.08 | 0.63 ± 0.18 | 2.05 ± 0.08 |
| Lysozyme | Guinier | 14.64 ± 0.05 | 14.28 ± 0.12 | 12.44 ± 0.52 | 0.36 ± 0.11 | 2.2 ± 0.52 |
| | $P(r)$ | 14.46 ± 0.01 | 14.66 ± 0.12 | 12.23 ± 0.10 | -0.2 ± 0.12 | 2.23 ± 0.1 |
| Urate Oxidase | Guinier | 32.30 ± 0.06 | 31.57 ± 0.42 | 31.18 ± 0.12 | 0.73 ± 0.42 | 1.02 ± 0.13 |
| | $P(r)$ | 31.63 ± 0.01 | 31.67 ± 0.23 | 30.84 ± 0.04 | -0.4 ± 0.23 | 0.79 ± 0.04 |

| force field | water model | $R_g^{Prot}$ (Å) | $R_g^{SAXS}$ (Å) | $R_g^{SANS\ H_2O}$ (Å) | $R_g^{SANS\ D_2O}$ (Å) | $R_g^{SAXS} - R_g^{Prot}$ (Å) | $R_g^{SANS\ H_2O} - R_g^{Prot}$ (Å) | $R_g^{SANS\ D_2O} - R_g^{Prot}$ (Å) | $R_g^{SAXS} - R_g^{SANS\ H_2O}$ (Å) | $R_g^{SAXS} - R_g^{SANS\ D_2O}$ (Å) |
|---|---|---|---|---|---|---|---|---|---|---|
| c36 | SPC/E | 14.69 ± 0.03 | 15.10 ± 0.07 | 14.67 ± 0.03 | 14.21 ± 0.04 | 0.42 ± 0.06 | -0.02 ± 0.02 | -0.47 ± 0.04 | 0.43 ± 0.06 | 0.89 ± 0.09 |
| ff14SB | SPC/E | 14.68 ± 0.03 | 15.36 ± 0.04 | 14.66 ± 0.03 | 13.99 ± 0.04 | 0.68 ± 0.03 | -0.02 ± 0.01 | -0.69 ± 0.03 | 0.70 ± 0.03 | 1.37 ± 0.05 |
| ff99SB-ildn | SPC/E | 14.51 ± 0.02 | 15.11 ± 0.06 | 14.50 ± 0.03 | 13.91 ± 0.05 | 0.60 ± 0.05 | -0.01 ± 0.02 | -0.60 ± 0.04 | 0.61 ± 0.05 | 1.20 ± 0.09 |
| c36 | cTIP3P | 14.65 ± 0.05 | 15.08 ± 0.06 | 14.67 ± 0.05 | 14.19 ± 0.04 | 0.43 ± 0.04 | 0.02 ± 0.02 | -0.46 ± 0.03 | 0.41 ± 0.04 | 0.89 ± 0.05 |
| c36 | TIP3P | 14.76 ± 0.03 | 15.28 ± 0.05 | 14.75 ± 0.03 | 14.21 ± 0.04 | 0.52 ± 0.04 | -0.01 ± 0.02 | -0.55 ± 0.04 | 0.53 ± 0.04 | 1.07 ± 0.06 |
| ff14SB | TIP3P | 14.75 ± 0.02 | 15.37 ± 0.07 | 14.72 ± 0.02 | 14.05 ± 0.05 | 0.62 ± 0.06 | -0.03 ± 0.01 | -0.70 ± 0.05 | 0.65 ± 0.06 | 1.32 ± 0.11 |
| ff99SB-ildn | TIP3P | 14.73 ± 0.02 | 15.36 ± 0.05 | 14.69 ± 0.02 | 14.04 ± 0.04 | 0.63 ± 0.05 | -0.04 ± 0.02 | -0.69 ± 0.04 | 0.67 ± 0.05 | 1.32 ± 0.08 |
| ff15fb | TIP3P-FB | 14.51 ± 0.02 | 15.12 ± 0.06 | 14.48 ± 0.02 | 13.86 ± 0.04 | 0.61 ± 0.05 | -0.03 ± 0.02 | -0.65 ± 0.05 | 0.64 ± 0.05 | 1.26 ± 0.09 |
| c36 | OPC3 | 14.55 ± 0.01 | 15.03 ± 0.05 | 14.53 ± 0.02 | 14.01 ± 0.05 | 0.48 ± 0.05 | -0.02 ± 0.02 | -0.53 ± 0.04 | 0.50 ± 0.05 | 1.01 ± 0.09 |
| ff99SBws | OPC3 | 14.74 ± 0.03 | 15.32 ± 0.04 | 14.70 ± 0.03 | 14.07 ± 0.03 | 0.58 ± 0.04 | -0.03 ± 0.02 | -0.66 ± 0.03 | 0.62 ± 0.04 | 1.25 ± 0.06 |
| c36 | OPC | 14.65 ± 0.02 | 15.33 ± 0.07 | 14.63 ± 0.02 | 13.95 ± 0.05 | 0.68 ± 0.06 | -0.02 ± 0.02 | -0.69 ± 0.06 | 0.70 ± 0.06 | 1.37 ± 0.11 |
| ff99SBws | OPC | 14.58 ± 0.04 | 14.98 ± 0.08 | 14.57 ± 0.03 | 14.15 ± 0.04 | 0.40 ± 0.06 | -0.01 ± 0.01 | -0.44 ± 0.04 | 0.41 ± 0.06 | 0.83 ± 0.10 |
| ff99SBws | TIP4P-D | 14.90 ± 0.06 | 15.44 ± 0.08 | 14.84 ± 0.06 | 14.25 ± 0.08 | 0.55 ± 0.06 | -0.05 ± 0.02 | -0.65 ± 0.05 | 0.60 ± 0.06 | 1.19 ± 0.11 |
| DES-amber | TIP4P-D | 14.65 ± 0.02 | 15.17 ± 0.05 | 14.60 ± 0.02 | 14.06 ± 0.03 | 0.52 ± 0.05 | -0.05 ± 0.02 | -0.59 ± 0.03 | 0.57 ± 0.04 | 1.11 ± 0.07 |
| DES-amberSF1.0 | TIP4P-D | 14.79 ± 0.03 | 15.41 ± 0.04 | 14.74 ± 0.02 | 14.09 ± 0.03 | 0.61 ± 0.04 | -0.05 ± 0.01 | -0.70 ± 0.02 | 0.67 ± 0.03 | 1.31 ± 0.05 |
| a99SBdisp | a99SBdisp-water | 14.72 ± 0.02 | 15.29 ± 0.06 | 14.70 ± 0.02 | 14.09 ± 0.04 | 0.57 ± 0.05 | -0.03 ± 0.01 | -0.63 ± 0.04 | 0.59 ± 0.05 | 1.20 ± 0.09 |
| ff99SBws | TIP4P/2005 | 14.75 ± 0.02 | 15.42 ± 0.05 | 14.72 ± 0.02 | 14.04 ± 0.05 | 0.66 ± 0.05 | -0.04 ± 0.02 | -0.71 ± 0.05 | 0.70 ± 0.05 | 1.37 ± 0.09 |
| ff99SBws | TIP4P/2005s | 14.78 ± 0.03 | 15.49 ± 0.05 | 14.73 ± 0.03 | 13.99 ± 0.04 | 0.71 ± 0.05 | -0.05 ± 0.02 | -0.79 ± 0.03 | 0.76 ± 0.04 | 1.50 ± 0.07 |

Table S3: $R_g$, $\Delta R_g$, and $\Delta R_g^{SAS}$ values from free MD simulations of 18 different combinations of protein force fields and water models for the protein RNaseA.

| force field | water model | $R_g^{Prot}$ (Å) | $R_g^{SAXS}$ (Å) | $R_g^{SANS\ H_2O}$ (Å) | $R_g^{SANS\ D_2O}$ (Å) | $R_g^{SAXS} - R_g^{Prot}$ (Å) | $R_g^{SANS\ H_2O} - R_g^{Prot}$ (Å) | $R_g^{SANS\ D_2O} - R_g^{Prot}$ (Å) | $R_g^{SAXS} - R_g^{SANS\ H_2O}$ (Å) | $R_g^{SAXS} - R_g^{SANS\ D_2O}$ (Å) |
|---|---|---|---|---|---|---|---|---|---|---|
| c36 | SPC/E | 14.17 ± 0.02 | 14.95 ± 0.04 | 14.32 ± 0.03 | 13.11 ± 0.04 | 0.78 ± 0.05 | 0.15 ± 0.02 | -1.06 ± 0.03 | 0.63 ± 0.05 | 1.84 ± 0.07 |
| ff14SB | SPC/E | 14.13 ± 0.01 | 14.78 ± 0.04 | 14.25 ± 0.02 | 13.11 ± 0.04 | 0.65 ± 0.04 | 0.12 ± 0.02 | -1.02 ± 0.04 | 0.53 ± 0.04 | 1.66 ± 0.07 |
| ff99SB-ildn | SPC/E | 14.06 ± 0.01 | 14.82 ± 0.04 | 14.18 ± 0.01 | 12.99 ± 0.04 | 0.76 ± 0.04 | 0.12 ± 0.01 | -1.07 ± 0.04 | 0.64 ± 0.05 | 1.83 ± 0.08 |
| c36 | cTIP3P | 14.31 ± 0.02 | 14.98 ± 0.03 | 14.43 ± 0.02 | 13.31 ± 0.03 | 0.66 ± 0.03 | 0.12 ± 0.01 | -1.00 ± 0.03 | 0.54 ± 0.03 | 1.66 ± 0.06 |
| c36 | TIP3P | 14.08 ± 0.02 | 14.80 ± 0.05 | 14.22 ± 0.03 | 13.11 ± 0.04 | 0.72 ± 0.04 | 0.14 ± 0.02 | -0.97 ± 0.04 | 0.58 ± 0.05 | 1.69 ± 0.07 |
| ff14SB | TIP3P | 14.12 ± 0.01 | 14.86 ± 0.04 | 14.25 ± 0.01 | 13.10 ± 0.03 | 0.74 ± 0.04 | 0.12 ± 0.01 | -1.02 ± 0.03 | 0.62 ± 0.04 | 1.76 ± 0.06 |
| ff99SB-ildn | TIP3P | 14.06 ± 0.02 | 14.84 ± 0.03 | 14.19 ± 0.03 | 13.02 ± 0.02 | 0.78 ± 0.03 | 0.12 ± 0.01 | -1.04 ± 0.02 | 0.65 ± 0.02 | 1.82 ± 0.03 |
| ff15fb | TIP3P-FB | 14.02 ± 0.01 | 14.73 ± 0.05 | 14.14 ± 0.02 | 12.90 ± 0.04 | 0.71 ± 0.05 | 0.12 ± 0.02 | -1.12 ± 0.04 | 0.59 ± 0.04 | 1.83 ± 0.08 |
| c36 | OPC3 | 14.13 ± 0.02 | 14.92 ± 0.04 | 14.25 ± 0.03 | 13.02 ± 0.05 | 0.78 ± 0.04 | 0.12 ± 0.02 | -1.11 ± 0.04 | 0.67 ± 0.05 | 1.89 ± 0.07 |
| ff99SBws | OPC3 | 14.01 ± 0.02 | 14.76 ± 0.04 | 14.11 ± 0.02 | 12.93 ± 0.04 | 0.76 ± 0.04 | 0.10 ± 0.01 | -1.08 ± 0.03 | 0.65 ± 0.03 | 1.84 ± 0.06 |
| c36 | OPC | 14.30 ± 0.03 | 14.94 ± 0.06 | 14.41 ± 0.03 | 13.29 ± 0.04 | 0.63 ± 0.05 | 0.11 ± 0.02 | -1.01 ± 0.05 | 0.52 ± 0.05 | 1.64 ± 0.09 |
| ff99SBws | OPC | 14.12 ± 0.01 | 14.82 ± 0.05 | 14.23 ± 0.02 | 13.06 ± 0.04 | 0.69 ± 0.05 | 0.10 ± 0.02 | -1.07 ± 0.04 | 0.59 ± 0.05 | 1.76 ± 0.09 |
| ff99SBws | TIP4P-D | 14.12 ± 0.02 | 14.84 ± 0.03 | 14.26 ± 0.02 | 13.05 ± 0.03 | 0.72 ± 0.03 | 0.14 ± 0.01 | -1.07 ± 0.03 | 0.58 ± 0.03 | 1.79 ± 0.06 |
| DES-amber | TIP4P-D | 14.25 ± 0.01 | 15.02 ± 0.04 | 14.37 ± 0.01 | 13.14 ± 0.03 | 0.77 ± 0.04 | 0.12 ± 0.01 | -1.12 ± 0.03 | 0.65 ± 0.04 | 1.89 ± 0.07 |
| DES-amberSF1.0 | TIP4P-D | 14.22 ± 0.02 | 15.06 ± 0.04 | 14.31 ± 0.02 | 12.99 ± 0.03 | 0.84 ± 0.04 | 0.10 ± 0.02 | -1.23 ± 0.03 | 0.74 ± 0.04 | 2.07 ± 0.06 |
| a99SBdisp | a99SBdisp-water | 14.10 ± 0.03 | 14.82 ± 0.06 | 14.24 ± 0.03 | 13.07 ± 0.04 | 0.71 ± 0.05 | 0.14 ± 0.01 | -1.04 ± 0.04 | 0.57 ± 0.05 | 1.75 ± 0.08 |
| ff99SBws | TIP4P/2005 | 13.97 ± 0.02 | 14.75 ± 0.04 | 14.09 ± 0.03 | 12.97 ± 0.03 | 0.78 ± 0.03 | 0.12 ± 0.01 | -1.00 ± 0.04 | 0.66 ± 0.03 | 1.78 ± 0.06 |
| ff99SBws | TIP4P/2005s | 14.03 ± 0.03 | 14.83 ± 0.05 | 14.13 ± 0.03 | 12.91 ± 0.04 | 0.80 ± 0.04 | 0.10 ± 0.01 | -1.12 ± 0.04 | 0.70 ± 0.04 | 1.92 ± 0.07 |

Table S4: $R_g$, $\Delta R_g$, and $\Delta R_g^{SAS}$ values from free MD simulations of 18 different combinations of protein force fields and water models for the protein xylanase.

| force field | water model | $R_g^{Prot}$ (Å) | $R_g^{SAXS}$ (Å) | $R_g^{SANS\ H_2O}$ (Å) | $R_g^{SANS\ D_2O}$ (Å) | $R_g^{SAXS} - R_g^{Prot}$ (Å) | $R_g^{SANS\ H_2O} - R_g^{Prot}$ (Å) | $R_g^{SANS\ D_2O} - R_g^{Prot}$ (Å) | $R_g^{SAXS} - R_g^{SANS\ H_2O}$ (Å) | $R_g^{SAXS} - R_g^{SANS\ D_2O}$ (Å) |
|---|---|---|---|---|---|---|---|---|---|---|
| c36 | SPC/E | 32.22 ± 0.01 | 34.00 ± 0.01 | 32.63 ± 0.01 | 31.48 ± 0.02 | 1.78 ± 0.02 | 0.41 ± 0.01 | -0.75 ± 0.02 | 1.37 ± 0.02 | 2.52 ± 0.03 |
| ff14SB | SPC/E | 32.15 ± 0.01 | 33.74 ± 0.02 | 32.59 ± 0.01 | 31.53 ± 0.02 | 1.59 ± 0.03 | 0.44 ± 0.01 | -0.63 ± 0.02 | 1.15 ± 0.02 | 2.21 ± 0.03 |
| ff99SB-ildn | SPC/E | 32.28 ± 0.01 | 33.85 ± 0.02 | 32.71 ± 0.01 | 31.67 ± 0.01 | 1.57 ± 0.02 | 0.43 ± 0.01 | -0.61 ± 0.01 | 1.14 ± 0.02 | 2.18 ± 0.02 |
| c36 | cTIP3P | 32.46 ± 0.02 | 34.08 ± 0.02 | 32.90 ± 0.02 | 31.82 ± 0.02 | 1.62 ± 0.01 | 0.43 ± 0.01 | -0.64 ± 0.01 | 1.19 ± 0.02 | 2.26 ± 0.02 |
| c36 | TIP3P | 32.26 ± 0.01 | 33.97 ± 0.02 | 32.67 ± 0.01 | 31.55 ± 0.02 | 1.71 ± 0.02 | 0.41 ± 0.01 | -0.71 ± 0.02 | 1.30 ± 0.02 | 2.43 ± 0.03 |
| ff14SB | TIP3P | 32.22 ± 0.01 | 33.73 ± 0.02 | 32.66 ± 0.01 | 31.66 ± 0.02 | 1.51 ± 0.02 | 0.45 ± 0.01 | -0.55 ± 0.02 | 1.07 ± 0.02 | 2.07 ± 0.03 |
| ff99SB-ildn | TIP3P | 32.39 ± 0.01 | 33.89 ± 0.01 | 32.84 ± 0.01 | 31.84 ± 0.02 | 1.49 ± 0.01 | 0.44 ± 0.01 | -0.55 ± 0.02 | 1.05 ± 0.02 | 2.04 ± 0.02 |
| ff15fb | TIP3P-FB | 32.06 ± 0.01 | 33.54 ± 0.02 | 32.48 ± 0.01 | 31.44 ± 0.01 | 1.47 ± 0.02 | 0.42 ± 0.01 | -0.62 ± 0.02 | 1.05 ± 0.02 | 2.10 ± 0.03 |
| c36 | OPC3 | 32.24 ± 0.01 | 33.93 ± 0.02 | 32.65 ± 0.01 | 31.57 ± 0.01 | 1.69 ± 0.02 | 0.42 ± 0.01 | -0.67 ± 0.01 | 1.27 ± 0.02 | 2.36 ± 0.03 |
| ff99SBws | OPC3 | 32.33 ± 0.01 | 33.93 ± 0.01 | 32.76 ± 0.01 | 31.70 ± 0.01 | 1.60 ± 0.01 | 0.44 ± 0.01 | -0.62 ± 0.01 | 1.16 ± 0.01 | 2.22 ± 0.02 |
| c36 | OPC | 32.30 ± 0.01 | 34.02 ± 0.03 | 32.71 ± 0.01 | 31.60 ± 0.02 | 1.72 ± 0.03 | 0.41 ± 0.01 | -0.70 ± 0.02 | 1.31 ± 0.03 | 2.42 ± 0.04 |
| ff99SBws | OPC | 32.44 ± 0.02 | 34.00 ± 0.02 | 32.87 ± 0.02 | 31.82 ± 0.03 | 1.56 ± 0.02 | 0.43 ± 0.01 | -0.62 ± 0.02 | 1.13 ± 0.02 | 2.18 ± 0.03 |
| ff99SBws | TIP4P-D | 32.42 ± 0.02 | 34.04 ± 0.02 | 32.85 ± 0.03 | 31.77 ± 0.03 | 1.62 ± 0.02 | 0.43 ± 0.01 | -0.65 ± 0.01 | 1.19 ± 0.02 | 2.27 ± 0.03 |
| DES-amber | TIP4P-D | 32.30 ± 0.01 | 33.77 ± 0.02 | 32.72 ± 0.01 | 31.78 ± 0.02 | 1.47 ± 0.02 | 0.42 ± 0.01 | -0.53 ± 0.01 | 1.05 ± 0.02 | 2.00 ± 0.03 |
| DES-amberSF1.0 | TIP4P-D | 32.35 ± 0.01 | 33.93 ± 0.02 | 32.78 ± 0.02 | 31.74 ± 0.02 | 1.58 ± 0.02 | 0.43 ± 0.01 | -0.60 ± 0.01 | 1.15 ± 0.02 | 2.19 ± 0.03 |
| a99SBdisp | a99SBdisp-water | 32.29 ± 0.02 | 33.83 ± 0.03 | 32.73 ± 0.02 | 31.66 ± 0.02 | 1.54 ± 0.02 | 0.44 ± 0.01 | -0.62 ± 0.02 | 1.10 ± 0.02 | 2.17 ± 0.03 |
| ff99SBws | TIP4P/2005 | 32.36 ± 0.02 | 34.01 ± 0.02 | 32.78 ± 0.02 | 31.69 ± 0.03 | 1.65 ± 0.02 | 0.43 ± 0.01 | -0.66 ± 0.02 | 1.23 ± 0.02 | 2.32 ± 0.03 |
| ff99SBws | TIP4P/2005s | 32.50 ± 0.03 | 34.16 ± 0.04 | 32.94 ± 0.03 | 31.84 ± 0.02 | 1.66 ± 0.02 | 0.44 ± 0.01 | -0.66 ± 0.01 | 1.21 ± 0.02 | 2.31 ± 0.03 |

Table S5: $R_g$, $\Delta R_g$, and $\Delta R_g^{SAS}$ values from free MD simulations of 18 different combinations of protein force fields and water models for the protein glucose isomerase.

| force field | water model | $R_g^{Prot}$ (Å) | $R_g^{SAXS}$ (Å) | $R_g^{SANS\ H_2O}$ (Å) | $R_g^{SANS\ D_2O}$ (Å) | $R_g^{SAXS} - R_g^{Prot}$ (Å) | $R_g^{SANS\ H_2O} - R_g^{Prot}$ (Å) | $R_g^{SANS\ D_2O} - R_g^{Prot}$ (Å) | $R_g^{SAXS} - R_g^{SANS\ H_2O}$ (Å) | $R_g^{SAXS} - R_g^{SANS\ D_2O}$ (Å) |
|---|---|---|---|---|---|---|---|---|---|---|
| c36 | SPC/E | 14.17 ± 0.02 | 14.95 ± 0.04 | 14.32 ± 0.03 | 13.11 ± 0.04 | 0.78 ± 0.05 | 0.15 ± 0.02 | -1.06 ± 0.03 | 0.63 ± 0.05 | 1.84 ± 0.07 |
| ff14SB | SPC/E | 14.13 ± 0.01 | 14.78 ± 0.04 | 14.25 ± 0.02 | 13.11 ± 0.04 | 0.65 ± 0.04 | 0.12 ± 0.02 | -1.02 ± 0.04 | 0.53 ± 0.04 | 1.66 ± 0.07 |
| ff99SB-ildn | SPC/E | 14.06 ± 0.01 | 14.82 ± 0.04 | 14.18 ± 0.01 | 12.99 ± 0.04 | 0.76 ± 0.04 | 0.12 ± 0.01 | -1.07 ± 0.04 | 0.64 ± 0.05 | 1.83 ± 0.08 |
| c36 | cTIP3P | 14.31 ± 0.02 | 14.98 ± 0.03 | 14.43 ± 0.02 | 13.31 ± 0.03 | 0.66 ± 0.03 | 0.12 ± 0.01 | -1.00 ± 0.03 | 0.54 ± 0.03 | 1.66 ± 0.06 |
| c36 | TIP3P | 14.08 ± 0.02 | 14.80 ± 0.05 | 14.22 ± 0.03 | 13.11 ± 0.04 | 0.72 ± 0.04 | 0.14 ± 0.02 | -0.97 ± 0.04 | 0.58 ± 0.05 | 1.69 ± 0.07 |
| ff14SB | TIP3P | 14.12 ± 0.01 | 14.86 ± 0.04 | 14.25 ± 0.01 | 13.10 ± 0.03 | 0.74 ± 0.04 | 0.12 ± 0.01 | -1.02 ± 0.03 | 0.62 ± 0.04 | 1.76 ± 0.06 |
| ff99SB-ildn | TIP3P | 14.06 ± 0.02 | 14.84 ± 0.03 | 14.19 ± 0.03 | 13.02 ± 0.02 | 0.78 ± 0.03 | 0.12 ± 0.01 | -1.04 ± 0.02 | 0.65 ± 0.02 | 1.82 ± 0.03 |
| ff15fb | TIP3P-FB | 14.02 ± 0.01 | 14.73 ± 0.05 | 14.14 ± 0.02 | 12.90 ± 0.04 | 0.71 ± 0.05 | 0.12 ± 0.02 | -1.12 ± 0.04 | 0.59 ± 0.04 | 1.83 ± 0.08 |
| c36 | OPC3 | 14.13 ± 0.02 | 14.92 ± 0.04 | 14.25 ± 0.03 | 13.02 ± 0.05 | 0.78 ± 0.04 | 0.12 ± 0.02 | -1.11 ± 0.04 | 0.67 ± 0.05 | 1.89 ± 0.07 |
| ff99SBws | OPC3 | 14.01 ± 0.02 | 14.76 ± 0.04 | 14.11 ± 0.02 | 12.93 ± 0.04 | 0.76 ± 0.04 | 0.10 ± 0.01 | -1.08 ± 0.03 | 0.65 ± 0.03 | 1.84 ± 0.06 |
| c36 | OPC | 14.30 ± 0.03 | 14.94 ± 0.06 | 14.41 ± 0.03 | 13.29 ± 0.04 | 0.63 ± 0.05 | 0.11 ± 0.02 | -1.01 ± 0.05 | 0.52 ± 0.05 | 1.64 ± 0.09 |
| ff99SBws | OPC | 14.12 ± 0.01 | 14.82 ± 0.05 | 14.23 ± 0.02 | 13.06 ± 0.04 | 0.69 ± 0.05 | 0.10 ± 0.02 | -1.07 ± 0.04 | 0.59 ± 0.05 | 1.76 ± 0.09 |
| ff99SBws | TIP4P-D | 14.12 ± 0.02 | 14.84 ± 0.03 | 14.26 ± 0.02 | 13.05 ± 0.03 | 0.72 ± 0.03 | 0.14 ± 0.01 | -1.07 ± 0.03 | 0.58 ± 0.03 | 1.79 ± 0.06 |
| DES-amber | TIP4P-D | 14.25 ± 0.01 | 15.02 ± 0.04 | 14.37 ± 0.01 | 13.14 ± 0.03 | 0.77 ± 0.04 | 0.12 ± 0.01 | -1.12 ± 0.03 | 0.65 ± 0.04 | 1.89 ± 0.07 |
| DES-amberSF1.0 | TIP4P-D | 14.22 ± 0.02 | 15.06 ± 0.04 | 14.31 ± 0.02 | 12.99 ± 0.03 | 0.84 ± 0.04 | 0.10 ± 0.02 | -1.23 ± 0.03 | 0.74 ± 0.04 | 2.07 ± 0.06 |
| a99SBdisp | a99SBdisp-water | 14.10 ± 0.03 | 14.82 ± 0.06 | 14.24 ± 0.03 | 13.07 ± 0.04 | 0.71 ± 0.05 | 0.14 ± 0.01 | -1.04 ± 0.04 | 0.57 ± 0.05 | 1.75 ± 0.08 |
| ff99SBws | TIP4P/2005 | 13.97 ± 0.02 | 14.75 ± 0.04 | 14.09 ± 0.03 | 12.97 ± 0.03 | 0.78 ± 0.03 | 0.12 ± 0.01 | -1.00 ± 0.04 | 0.66 ± 0.03 | 1.78 ± 0.06 |
| ff99SBws | TIP4P/2005s | 14.03 ± 0.03 | 14.83 ± 0.05 | 14.13 ± 0.03 | 12.91 ± 0.04 | 0.80 ± 0.04 | 0.10 ± 0.01 | -1.12 ± 0.04 | 0.70 ± 0.04 | 1.92 ± 0.07 |

Table S6: $R_g$, $\Delta R_g$, and $\Delta R_g^{SAS}$ values from free MD simulations of 18 different combinations of protein force fields and water models for the protein lysozyme.

| force field | water model | $R_g^{Prot}$ (Å) | $R_g^{SAXS}$ (Å) | $R_g^{SANS\ H_2O}$ (Å) | $R_g^{SANS\ D_2O}$ (Å) | $R_g^{SAXS} - R_g^{Prot}$ (Å) | $R_g^{SANS\ H_2O} - R_g^{Prot}$ (Å) | $R_g^{SANS\ D_2O} - R_g^{Prot}$ (Å) | $R_g^{SAXS} - R_g^{SANS\ H_2O}$ (Å) | $R_g^{SAXS} - R_g^{SANS\ D_2O}$ (Å) |
|---|---|---|---|---|---|---|---|---|---|---|
| c36 | SPC/E | 31.19 ± 0.01 | 32.15 ± 0.02 | 31.57 ± 0.01 | 31.22 ± 0.02 | 0.96 ± 0.02 | 0.38 ± 0.01 | 0.03 ± 0.02 | 0.58 ± 0.02 | 0.92 ± 0.03 |
| ff14SB | SPC/E | 31.27 ± 0.02 | 32.28 ± 0.03 | 31.65 ± 0.01 | 31.25 ± 0.02 | 1.02 ± 0.03 | 0.38 ± 0.01 | -0.02 ± 0.02 | 0.63 ± 0.03 | 1.04 ± 0.04 |
| ff99SB-ildn | SPC/E | 31.50 ± 0.02 | 32.58 ± 0.03 | 31.88 ± 0.02 | 31.46 ± 0.02 | 1.08 ± 0.02 | 0.37 ± 0.01 | -0.05 ± 0.01 | 0.70 ± 0.02 | 1.12 ± 0.03 |
| c36 | cTIP3P | 31.30 ± 0.02 | 32.14 ± 0.05 | 31.65 ± 0.02 | 31.39 ± 0.03 | 0.84 ± 0.04 | 0.35 ± 0.01 | 0.09 ± 0.02 | 0.49 ± 0.04 | 0.75 ± 0.05 |
| c36 | TIP3P | 31.23 ± 0.02 | 32.10 ± 0.02 | 31.59 ± 0.01 | 31.30 ± 0.02 | 0.87 ± 0.02 | 0.36 ± 0.01 | 0.07 ± 0.01 | 0.50 ± 0.02 | 0.80 ± 0.02 |
| ff14SB | TIP3P | 31.16 ± 0.02 | 32.14 ± 0.02 | 31.55 ± 0.02 | 31.16 ± 0.03 | 0.99 ± 0.02 | 0.39 ± 0.01 | -0.00 ± 0.01 | 0.60 ± 0.02 | 0.99 ± 0.02 |
| ff99SB-ildn | TIP3P | 31.31 ± 0.01 | 32.30 ± 0.02 | 31.69 ± 0.01 | 31.31 ± 0.02 | 0.99 ± 0.02 | 0.39 ± 0.01 | 0.01 ± 0.02 | 0.60 ± 0.02 | 0.98 ± 0.03 |
| ff15fb | TIP3P-FB | 31.10 ± 0.02 | 32.01 ± 0.02 | 31.48 ± 0.02 | 31.11 ± 0.02 | 0.91 ± 0.02 | 0.38 ± 0.01 | 0.01 ± 0.01 | 0.53 ± 0.02 | 0.90 ± 0.02 |
| c36 | OPC3 | 31.21 ± 0.03 | 32.21 ± 0.05 | 31.58 ± 0.03 | 31.20 ± 0.03 | 1.00 ± 0.03 | 0.37 ± 0.01 | -0.01 ± 0.02 | 0.63 ± 0.03 | 1.01 ± 0.04 |
| ff99SBws | OPC3 | 31.49 ± 0.02 | 32.56 ± 0.02 | 31.86 ± 0.02 | 31.45 ± 0.03 | 1.07 ± 0.02 | 0.37 ± 0.01 | -0.04 ± 0.02 | 0.70 ± 0.02 | 1.11 ± 0.03 |
| c36 | OPC | 31.35 ± 0.02 | 32.28 ± 0.03 | 31.69 ± 0.02 | 31.39 ± 0.03 | 0.93 ± 0.02 | 0.34 ± 0.01 | 0.04 ± 0.02 | 0.59 ± 0.02 | 0.90 ± 0.03 |
| ff99SBws | OPC | 31.63 ± 0.03 | 32.63 ± 0.05 | 31.97 ± 0.02 | 31.64 ± 0.03 | 1.00 ± 0.04 | 0.35 ± 0.01 | 0.01 ± 0.02 | 0.65 ± 0.04 | 0.99 ± 0.05 |
| ff99SBws | TIP4P-D | 31.54 ± 0.02 | 32.56 ± 0.02 | 31.93 ± 0.02 | 31.54 ± 0.02 | 1.02 ± 0.02 | 0.39 ± 0.01 | -0.00 ± 0.01 | 0.63 ± 0.02 | 1.02 ± 0.03 |
| DES-amber | TIP4P-D | 31.35 ± 0.01 | 32.28 ± 0.03 | 31.70 ± 0.02 | 31.39 ± 0.02 | 0.94 ± 0.02 | 0.35 ± 0.01 | 0.04 ± 0.01 | 0.58 ± 0.02 | 0.89 ± 0.03 |
| DES-amberSF1.0 | TIP4P-D | 31.32 ± 0.02 | 32.40 ± 0.03 | 31.68 ± 0.02 | 31.26 ± 0.03 | 1.08 ± 0.02 | 0.36 ± 0.01 | -0.06 ± 0.01 | 0.72 ± 0.02 | 1.14 ± 0.03 |
| a99SBdisp | a99SBdisp-water | 31.38 ± 0.02 | 32.31 ± 0.03 | 31.76 ± 0.01 | 31.42 ± 0.01 | 0.93 ± 0.02 | 0.38 ± 0.01 | 0.03 ± 0.02 | 0.55 ± 0.03 | 0.89 ± 0.04 |
| ff99SBws | TIP4P/2005 | 31.47 ± 0.01 | 32.50 ± 0.02 | 31.84 ± 0.01 | 31.44 ± 0.01 | 1.03 ± 0.02 | 0.37 ± 0.01 | -0.03 ± 0.01 | 0.66 ± 0.02 | 1.06 ± 0.03 |
| ff99SBws | TIP4P/2005s | 31.57 ± 0.01 | 32.66 ± 0.02 | 31.94 ± 0.02 | 31.48 ± 0.02 | 1.09 ± 0.02 | 0.37 ± 0.01 | -0.09 ± 0.02 | 0.72 ± 0.03 | 1.18 ± 0.04 |

Table S7: $R_g$, $\Delta R_g$, and $\Delta R_g^{SAS}$ values from free MD simulations of 18 different combinations of protein force fields and water models for the protein urate oxidase.

(17) Jorgensen, W. L.; Chandrasekhar, J.; Madura, J. D.; Impey, R. W.; Klein, M. L.

Comparison of simple potential functions for simulating liquid water. *J. Chem. Phys.* **1983**, *79*, 926–935.
